# First superferromagnetic remanence characterization and scan optimization for super-resolution Magnetic Particle Imaging

**DOI:** 10.1101/2022.11.08.515719

**Authors:** K. L. Barry Fung, Caylin Colson, Jacob Bryan, Chinmoy Saayujya, Javier Mokkarala-Lopez, Allison Hartley, Khadija Yousuf, Renesmee Kuo, Yao Lu, Benjamin D. Fellows, Prashant Chandrasekharan, Steven M. Conolly

## Abstract

Magnetic particle imaging (MPI) is a sensitive, high contrast tracer modality that images superparamagnetic iron oxide nanoparticles (SPIOs), enabling radiation-free theranostic imaging. MPI resolution is currently limited by scanner and particle constraints. Recent tracers have experimentally shown 10x resolution and signal improvements, with dramatically sharper M-H curves. Experiments suggest that this results from interparticle interactions, conforming to literature definitions of superferromagnetism. We thus call our tracers superferromagnetic iron oxide nanoparticles (SFMIOs). While SFMIOs provide excellent signal and resolution, they exhibit hysteresis, with non-negligible remanence and coercivity. We provide the first report on MPI scanning with remanence and coercivity, including the first quantitative measurements of SFMIO remanence decay and reformation using a novel multi-echo pulse sequence. We also describe an SNR-optimized pulse sequence for SFMIOs under human electromagnetic safety limitations. The resolution from SFMIOs could enable clinical MPI with 10× reduced scanner selection fields, reducing hardware costs by up to 100×.

## Main Text

Magnetic particle imaging (MPI) is an emerging tracer modality^1,2^ that directly images the magnetization of superparamagnetic iron oxide nanoparticles (SPIOs) with positive, linear contrast. First described by Gleich and Weizenecker in 2005,^1^ MPI leverages the non-linear magnetic response of SPIOs to localise the SPIOs and generate an image proportional to tracer concentration.

Iron oxide nanoparticle tracers were first introduced to magnetic resonance imaging (MRI) by Lauterbur’s group in 1986.^3^ Superparamagnetic iron oxide nanoparticles (SPIOs) have been traditionally used in MRI, primarily as a T2*-weighted contrast agent. High concentrations of SPIOs produce dark signals on T2*-MRI, which radiologists term “negative contrast,” and different SPIO formulations can yield varied targeting and contrast.^4^ Unfortunately, it is challenging to distinguish dark T2* signal from a naturally dark signal (e.g., the MRI signal in the lungs, bones, cartilage). Hence, quantitative measurements with T2*-SPIOs remains challenging. ^5–9^ An exception here is liver MRI, where SPIOs are preferentially taken up by healthy liver tissue, leaving dysfunctional tissue (e.g., function lost due to liver cancer) with bright signal.^10^

In comparison, the MPI signal depends on the SPIO’s non-linear magnetic response. Unlike human tissue, which remains linear^7^ well above 3 T, SPIOs reach a saturation magnetization at low applied fields^11^ (typically ≈ 6 mT). This non-linear saturation yields the harmonics in MPI signals,^12^ and allows for positive-contrast detection of SPIOs with no background signal from tissues.^13^ Moreover, the MPI signal is robust to magnetic field inhomogeneities (5% field variations are well tolerated^14^ vs 10 ppm for MRI), and has minimal attenuation from tissue depth given its relatively low frequencies (max(*f*_sig_) ≪ 10 MHz),^13^ making MPI suitable for imaging the entire body. Given these upsides, MPI shows promise in imaging applications previously dominated by nuclear medicine, like pulmonary embolism detection,^15–17^ cancer,^18^ gut bleed,^19^ and white blood cell (neutrophil) imaging of cancer,^20^ infection and bone marrow function. ^21^ MPI also shows promise in theranostics, assisting in targeted drug delivery,^22^ cell therapy monitoring,^23,24^ and magnetic hyperthermia treatment. ^25–27^

However, despite its excellent safety profile and proven *in vivo* applications, MPI still faces a hurdle to clinical translation due to its resolution. MPI’s resolution is roughly 1 mm^12,28^ in preclinical scanners with intense 6.3 to 7T/m selection fields.^19,29,30^ This is not competitive as compared to 500 μm or smaller in other preclinical tracer modalities.^31^ While these resolutions are comparable, scaling up magnetic systems is incredibly expensive, with human versions of preclinical MPI scanners projected to require field strengths equivalent to multi-million dollar 7T MRIs. This would be an *order* of magnitude more expensive than positron emission tomography and X-ray computed tomography scanners. As such, MPI is in need of fundamental (and cost-effective) improvements to resolution.

Improving MPI’s fundamental resolution requires improving the SPIO’s magnetic response or the scanner’s gradient strength. Since MPI utilises inductive measurements of magnetization, its point spread function (PSF) is proportional to the derivative of the Langevin function,^2^ convolved with the SPIO’s magnetic relaxation behaviour^32^ in time. The resulting resolution, measured as full-width at half-max (FWHM), is approximately:

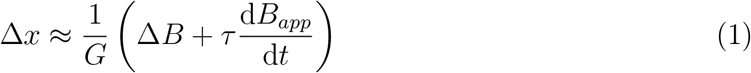

where Δ*B* is the width of the Langevin’s magnetic transition in T, *τ* is the magnetic relaxation time constant in s, d*B_app_*/d*t* is the slew rate of the magnetic scanner (in T/s), and *G* is the gradient in T/m. In MPI, the relaxation of a magnetic particle can be modeled as an exponential decay with time constant *τ*.^32^ Based on the scanner’s magnetic slew rate, the ideal magnetic resolution ΔB is thus blurred by the slew rate multiplied by signal decay constant (i.e., *τ*), as in Eq. 1.

Of the two approaches to improving MPI resolution, scaling gradient strength is costly and can require superconducting magnets costing millions.^33^ As a result, recent work has focused on modifying SPIO behaviour. Notably, the magnetic transition scales inversely with magnetic volume (i.e., 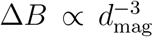), while the relaxation time scales with hydrodynamic volume and to the natural exponent of the magnetic volume 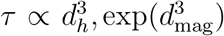.^34^ The net effect results in a reduction of sensitivity and resolution of the MPI system above a magnetic particle diameter threshold (dubbed the “relaxation wall”^28^). Empirically, this threshold was found to be 25nm for single core, magnetite particles (at 20mT amplitude, 20 kHz excitation). However, optimal tracers still only had effective resolutions of ≈ 1 mm.^12,28^ Strategies modifying acquisition waveforms ^25,35,36^ and reconstruction methods ^37,38^ have provided methods for bypassing the relaxation wall resolution limit, but each method suffers from long scan times, lower sensitivity, high SNR requirements, or limiting algorithmic priors.

Recently, we described high resolution, high sensitivity SPIOs^39^ that exhibited optimal behaviour at high concentrations and excitation amplitudes (Figure 1A-C, imaged in a 6.3T/m FFL scanner^19^). In MPI scanners, these SPIOs (Figure 1C-D) showed 20-fold higher resolution than shown in powder DC magnetometry (Fig. 1E). Briefly, these SPIOs are thought to apply fields on neighbouring SPIOs at higher concentrations (i.e., lower interparticle distances), effectively amplifying externally applied fields. This amplification increases as interparticle distance approaches one diameter, where the particles form chainlike mesostructures (Figure 1G).^40,41^ Importantly, a simple, positive-feedback model using a Langevin operator yields a very compelling magnetic model for these SPIOs (Figure 1H), predicting the observed hysteresis and PSF (Fig. 1I). This ensemble regenerative magnetic response has previously been described in literature as superferromagnetism^42–44^, but had not been examined in the context of inductive detection and magnetic particle imaging. We thus described these particles as superferromagnetic iron oxide nanoparticles (SFMIOs).

**Figure 1:**
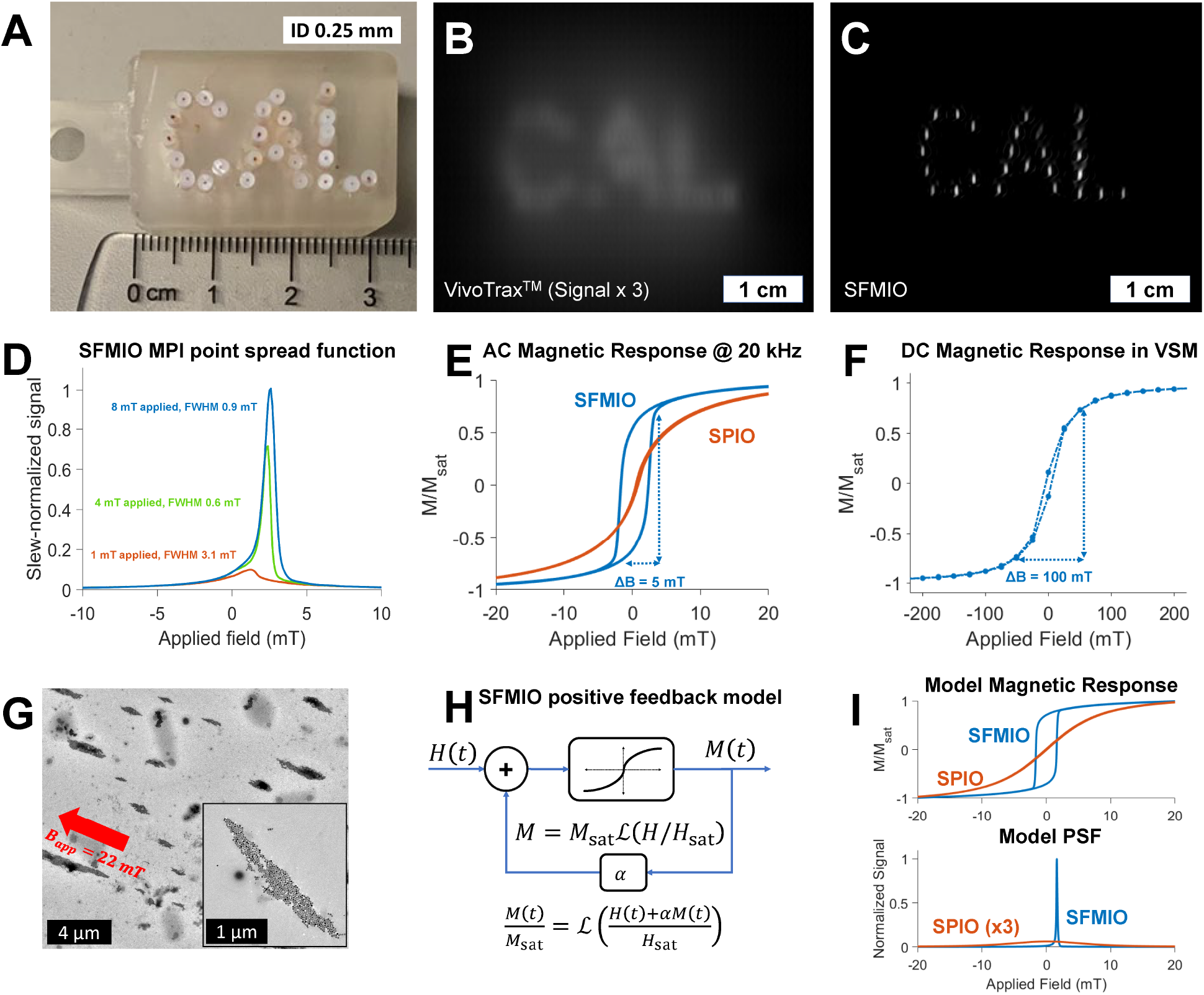
SFMIOs show 10-fold resolution in MPI: Recent single-core magnetite iron oxide nanoparticles, dubbed superferromagnetic iron oxide particles (SFMIOs) show 10-fold resolution improvement compared to ferrucarbotran (A-C). When these particles are at high concentration and are excited with a strong magnetic field, their peak signal and resolution greatly improves (D). These particles show strong hysteresis and sharp magnetic transitions (E) in MPI scanners in solution, but dried samples show superparamagnetic behaviour (F). This resolution improvement is hypothesized to be due to particle-particle interactions, which are possible thanks to the size of the particles (25 –30nm) vs. their coating (≈ 1 nm). These particles appear to form chain like structures when dried under a field (visualized under electron microscopy) (G, adapted with permission from Tay et al. 2021^39^). Indeed, a simple positive-feedback model (H), using the Langevin function as the system function, yields an magnetic response (I) that shows nearly identical behaviour, lending credence to the hypothesis.

With the resolution improvements offered by SFMIOs, scanner field strengths could be 10× weaker while maintaining MPI’s current resolution in humans, allowing for up to a 100× reduction in cost. However, while SFMIOs offer incredible benefits, its properties are highly dependent on the magnetically-generated SFMIO chains^39^ (Fig. 1G), which have time-varying hysteresis, remanence and coercivity. Assessing SFMIO characteristics to ensure safe and efficient SFMIO scans is thus essential for clinical translation for MPI. In this work, we characterize SFMIO remanence, its decay and reformation after magnetic polarization, and propose future MPI scan strategies to optimally and safely image SFMIOs.

### Results and Discussion

≈28-nm magnetite nanoparticles (SFMIOs) were synthesized via thermal decomposition^45^ and suspended in hexane, with a thin oleic acid coating. 40 μL of SFMIOs at 2.76mg Fe/mL (quantified by Perl’s Prussian Blue reaction) were measured in an arbitrary-waveform relaxometer.^46^ The point spread function (PSF) was measured using sinusoidal 20 kHz fields of 1-10mT amplitudes (*B*_Tx_), to assess their resolution, coercivity, and signal. Notably, the SFMIOs demonstrated super-resolution behavior for *B*_Tx_ ≥ *B*_th_, where *B*_th_ varied from 4mT to 8mT depending on the characteristics of the synthesized batch of particles.

Standard MPI sequences utilise purely sinusoidal waveforms, and can only show remanence evolution mixed with the effects of applied fields. As such, a novel magnetic pulse sequence (reminiscent of pulsed MPI^25^). was devised to measure magnetic remanence, utilising a home-built arbitrary-waveform relaxometer (Fig. 2A). ^46^ We describe the measurement signal of remanence decay phenomenologically in Eq. 2:

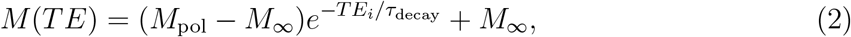

where *M* is the magnetization at time TE, *M*_pol_ is the magnetization of the fully polarized SPIO structure, *TE* is the echo time between pulses, *τ*_decay_ is the measured remanence decay constant, and *M*_∞_ is the steady-state magnetization, including any steady-state remanence.

**Figure 2:**
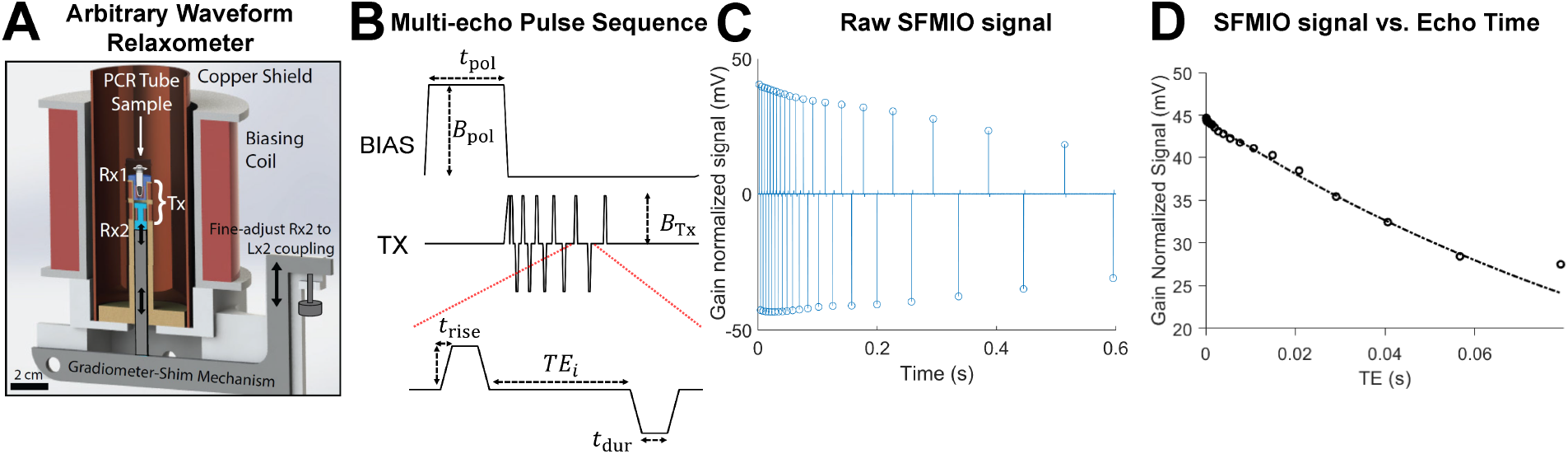
Measuring SFMIO remanence and remanence evolution: T measure the relatively slow variation in SFMIO remanence, we utilised an arbitrary wave relaxometer (A, reproduced from Tay et al. 2016^46^ with permission) to apply an arbitrary magnetic pulse sequence with minimal feedthrough. The pulse sequence (B) first polarizes the SFMIOs to form chains with a large (*B*_pol_) and prolonged (*t*_pol_) magnetic pulse, using a high-inductance, high efficiency bias coil. It then reads out remanence decay by applying faster trapezoidal pulses (defined by *t*_rise_, *t*_dur_, *B*_Tx_) with increasing interpulse spacing, dubbed echo times (*TE_i_*), to perform a multi-echo readout - the first-of-its-kind in MPI. This latter set of magnetic pulses is performed with a responsive, lower inductance transmit (Tx) coil, and allows inductive readout of slower decay constants. The raw signal (C) can then be used to extract a function of SFMIO signal versus TE as a surrogate for SFMIO remanence (D), which yielded a remanence decay constant at zero field of *τ*_decay_(*B*)|_*B*=0_ ≈120 ms.

The first-of-its-kind MPI multi-echo sequence (Fig. 2B) concatenates consecutive magnetic pulses with increasing echo time (TE) at 0 mT, in order to measure the effective remanence decay behaviour after polarization. Specifically, SFMIOs were polarized using strong fields (*B*_pol_ = 30 mT, *t*_pol_ = 30 s), and then measured with alternating trapezoidal pulses (*t*_rise_ = 10 μs, *t*_dur_ = 3 ms, *B*_Tx_ = ±32 mT) with increasing inter-pulse duration (*TE_i_* = [100 μs, 80 ms]). The resultant signal (Fig. 2C-D) represents remanence with increasing echo time *TE_i_*, providing a surrogate measurement of SFMIO remanence and its evolution during an MPI scan. Notably, this multi-echo approach allows for inductive measurement of slow decay constants that provide minimal signal.

As shown in Fig. 2D, the MPI signal of SFMIOs exponentially decayed with increasing TE at 0mT field (*τ*_decay_ ≈ 120 ms), but did not fully lose super-resolution behaviour. While the observed decay did not disrupt super-resolution behaviour in standard scans, any variation in amplitude could confound intraimage analysis.

To dissect the effects of our multi-echo sequence on SFMIO remanence, we examine its individual components. The prototypical sequence (shown in Fig 3A, top) traverses the SFMIO hysteresis curve (Fig 3A, bottom), first polarizing the sample (Fig 3A, i), then allowing the sample to sit at 0 mT for some echo time TE (Fig 3A, ii-iii), then measuring the resultant remanence and consequent transition *M*(*TE*) = *M*_TE_ + *M*_sat_ (Fig 3A, iv) as the sample repolarizes in the opposite direction.

**Figure 3:**
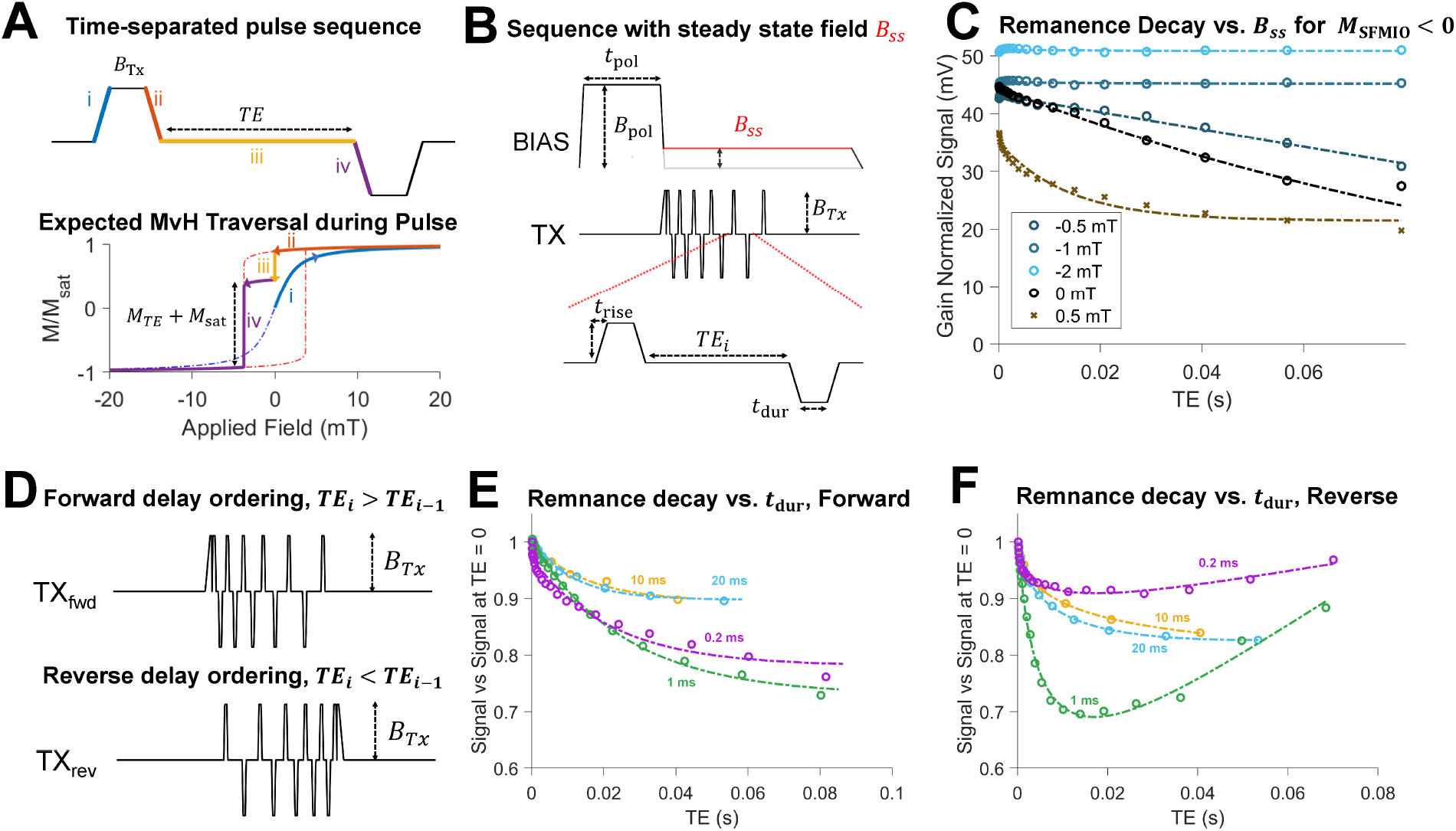
Characterizing remanence decay in MPI scans and chain reformation: Our time separated magnetic pulses (A, top) allow us to characterize the behaviour of SFMIO remanence over time. Specifically, by polarizing the particles with a pulse of amplitude *B*_Tx_ (i, ii), and allowing the sample to sit at zero field for some specified echo time TE (iii), we can read out using a secondary pulse to measure an MPI signal (iv), while also simultaneously repolarizing the particles. If we examine the traversal of the magnetic response curve (A, bottom), it is clear that this signal is a function of the remaining magnetization *M*_TE_ + *M*_sat_. (B) By modifying the multi-pulse sequence to have a steady-state field *B_SS_* during acquisition, we altered the field at which remanence decay (A, iii) occurs. The decay mechanics at various fields is shown in (C). Notably, for a chain polarized in the negative direction (*M*_SPIO_ < 0, pre-positive slew), small, anti-parallel fields showed the acceleration of decay *B_SS_* > 0, yellow), while parallel fields (*B_SS_* < 0, blue) showed reinforcement and decay suppression. Similar relationships were observed for chains polarized in the opposite directions. By modifying the delay order (D), we could see the effect of echo-ordering, and provide an estimate for the time constant for reformation after decay. For pulse lengths less than 10 ms, echo ordering produced wildly different decay patterns (E-F), while pulse lengths equal to or longer than 10 ms showed similar patterns (with lines as guides for the eyes). This leads to an estimate for 5*τ*_reformation_ = 10 ms, assuming that 5 time constants are sufficient for steady-state behaviour.

To further characterize remanence evolution, various pulse sequence parameters were modified, and the resultant decay pattern measured. To assess decay during non-zero fields, the sample was measured while applying various steady-state fields (*B_SS_* = [−4 mT, 1 mT]) (Fig. 3B). Fig. 3C shows the remanence evolution as a function of steady-state field for SFMIOs polarized in the negative direction (*M*_SFMIO_ < 0). When fields parallel to the structure (*B_SS_* < 0) were applied, the SFMIO signal showed little to no decay. In comparison, applying minimal anti-parallel fields (*B_SS_* > 0) showed greatly accelerated decay with (*τ*_decay_ ≈ 13 ms) even at *B_SS_* = 0.5 mT.

In the excitation field in a standard MPI scan, SFMIO chains will mostly tend to experience reinforcing fields, with anti-parallel fields occurring as the field sweeps past 0 mT to the coercive threshold opposing the SFMIO chain (*B*_coercivity_ ≈ ±4mT in Fig. 3). Thus, the minimum scan speed for SFMIOs should be governed by the transition through this region. Given that the fastest decay was shorter than the sampling period (for *B_SS_* approaching *B_coercivity_*, in Supplemental Fig. 2), the frequency threshold is only lower bound by our measurement (i.e., d*B*/d*t* > *B_coercivity_*/min(*τ_decay,meas_*) = 15 Hz). More generally, the minimum scan rate will be:

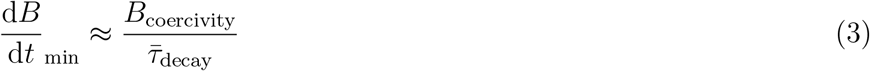

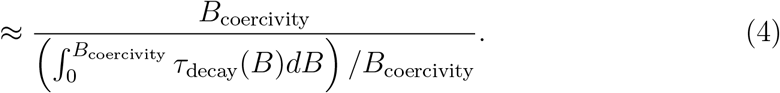

assuming a constant slew rate. This accounts for all decay rates between 0 mT and *B*_coercivity_. The derived lower bound is congruent with previous SFMIO measurements versus frequency, which showed super-resolution behaviour above 100 Hz (*B*_app_ = 20 mT).^39^ sets limits for SFMIO scan trajectory design.

To assess how quickly the SFMIO chains are reformed after perturbation, the echo ordering of the sequence was reversed (Fig. 3D), and the pulse duration *t*_dur_ was varied from *t*_dur_ = [1 ms − 20 ms]. The resultant signal is then a recursive composition of remanence decay and reformation,

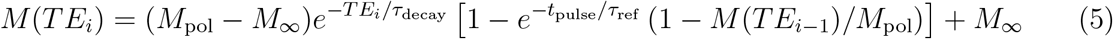

where *τ*_ref_ is a chain reformation constant and *M*(*TE*_*i*−1_) represents the previous echo’s signal for a given ordering. From this equation, pulse durations that are insufficient for full reformation (i.e., *t*_pulse_ faster than *τ*_ref_) should yield inconsistent measurements for varying echo orders, as contributions from initial measurements will appear in later echoes. However, for *t*_pulse_ ≫ *τ*_ref_, the term exp (–*t*_pulse_/*τ*_ref_) (representing contributions from previous echoes) nears zero, and Eq. 5 simplifies to Eq. 2.

The resultant decay patterns are shown in Figs 3E-F: for *t*_dur_ < 10 ms, the reverse sequences yield different measurements than forward sequences, visible as non-monotonic function of TE. In comparison, the decay patterns with consistent decay irrespective of delay order in Fig. 3 imply full chain reformation for *t*_dur_ ≥ 10 ms. This yields an estimate of *τ*_reformation_ ≈ 2 ms, assuming 5*τ*_reformation_ yields complete chain reformation.

Remanence decay and reformation can occur for various reasons. For one, the SFMIO chain could be an unstable colloid, dispersing at 0 mT and resulting in superparamagnetism. Reformation would then be the complete reformation of SFMIO chains. However, this seems unlikely based on the potential energies for interacting particles. Consider two SPIOs with aligned domains (Fig. 4A and B, top), under some magnetic field. Traditional DLVO models^47,48^ for charge-separated ferrofluids^49,50^ have two stable energy minima that can exist for these SPIOs (characteristic curve in Fig. 4A, bottom), due to electrostatic repulsion, Van der Waals interactions, and magnetic attraction. Colloidal dispersion can then occur when the magnetic field is removed and the secondary minima disappears.

**Figure 4:**
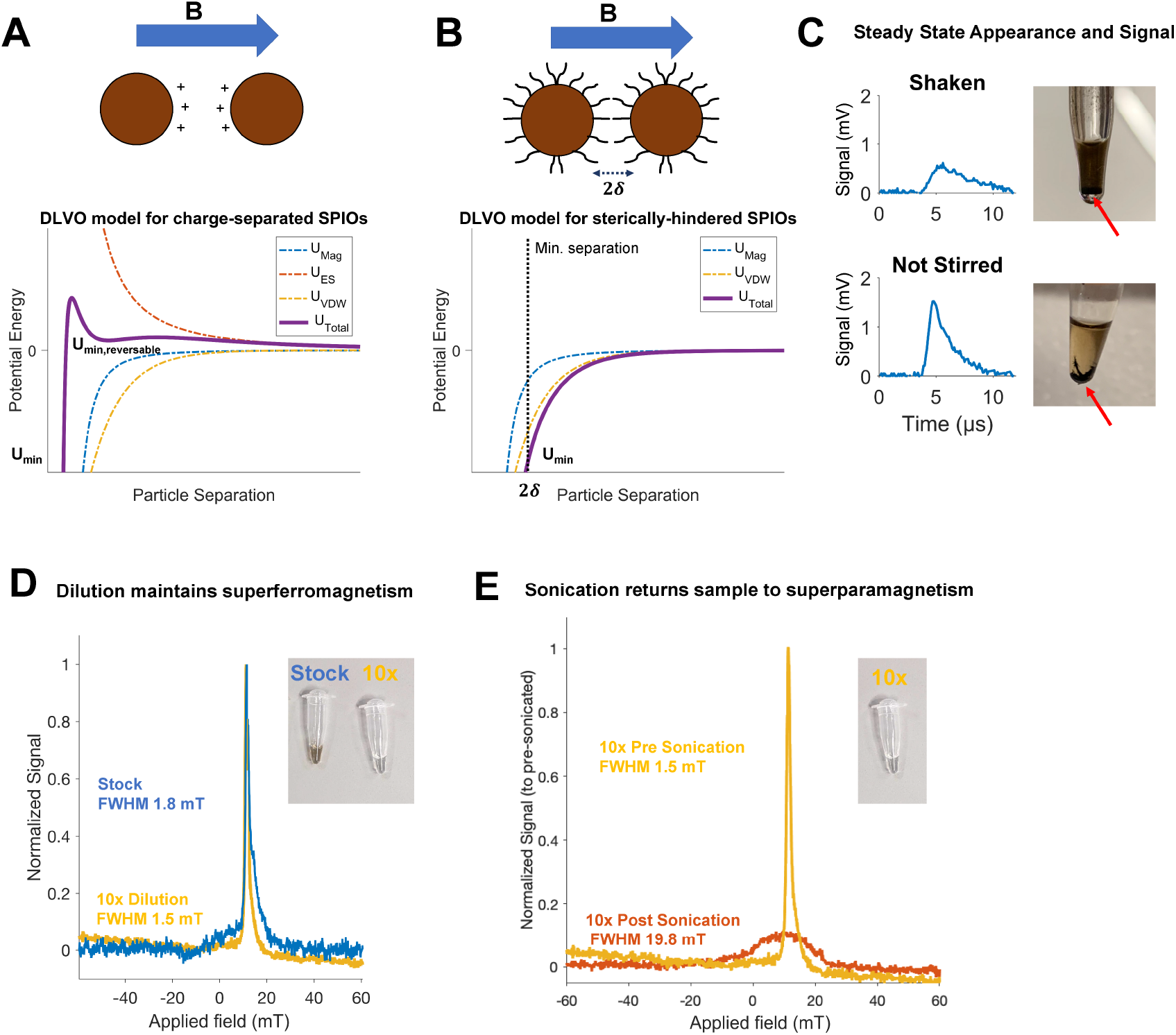
Potential mechanisms for SFMIO remanence decay: Models of colloid stability provide insight into SFMIO remanence. While traditional ferrofluid theory has shown spontaneous dissolution, this was for charge-separated particles in aqueous media. For these cases, DLVO theory^47–49^ (A) states that the potential for electrostatic repulsion, magnetic attraction, and Van der Waals attraction governs colloidal stability. Specifically, the attractive and repulsive forces can allow for unstable aggregation in charge-separated SPIOs in specific configurations. However, sterically-hindered SPIOs in non-polar media (as in this work) show a single, irreversible minima caused by both attractive forces (B). As such, it is unlikely that spontaneous dissolution of SFMIO chains is occurring. At steady state (*t* = 12 s, *t* ≫ 5*τ*_decay_) with manual agitation for 30s, the sample showed standard SPIO behaviour magnetically (C, top), with the SFMIOs visibly settling (as shown by the arrow). Conversely, for samples that were not shaken (or stirred) at steady-state (C, bottom), visual and magnetic inspection showed persistent chains and super-resolution activity. This was further confirmed by dilution of polarized SFMIOs, which maintained their super-resolution behavior (D), until after sonication (E).

However, for sterically-hindered particles in non-polar media (as in the experiments above), the electrostatic force from surface charge is insignificant and no secondary minima exists (Fig. 4B, bottom). Steric hindrance prevents bulk matter formation at ≈ 2*δ*, where *δ* is the surface ligands length.^51^ In this case, colloidal dispersion is not energetically favourable. Theoretically, applying an opposing field equivalent to SFMIO coercivity could negate magnetic attraction momentarily, leaving only Van der Waals interactions to stabilize the colloid. However, steady-state experiments (Fig. 3C) showed that even small anti-parallel fields (well below the coercivity) accelerate decay. Thus, the chains are likely not spontaneously breaking apart.

Instead, the SFMIO chains may simply be misaligned from the measurement axis. If left alone, individual particles may stick together, but the chains could point in different directions. ^41^ Previous work^39^ has shown that SFMIO super-resolution behaviour occurs only when the structure aligns with the measurement axis - if the chain is orthogonal to the measurement axis, the SFMIOs appear superparamagnetic. Similar hysteresis changes have been observed in cryo-magnetometry of bionized nanoferrite.^52^ A misalignment of the chains could allow for superferromagnetism, but with a reduced signal. Indeed, when the sample was measured after TE = 12 s (i.e., ≫ 5*τ*_decay_), we continued to observe super-resolution behaviour (Fig. 4C, bottom), and visually see SFMIO chaining. To ensure that the SFMIO chain was yielding steady-state superferromagnetism, the sample was mechanically agitated for 30 seconds at the same time point (TE = 12 s). As shown in (Fig 4C, top), only superparamagnetic behavior remains. This hypothesis also accounts for decay acceleration in small, opposing fields, as anti-parallel chains in a unidirectional field are in an unstable equilibrium, and any perturbation would result in magnetic torquing. ^53^

We further tested this hypothesis by examining polarized SFMIOs after dilution and sonication. Even after 10-fold dilution, the SFMIOs maintained superferromagnetism (Fig.4D). If spontaneous dissolution was occurring, this diluted solution should not show superferromagnetic behavior, as the sample is below the concentration threshold for superferromagnetism. ^39^ The observation otherwise suggests that the chains are maintained even through dilution. In comparison, sonication (Fig. 4E) was able to return the sample to superparamagnetism. As sonication imparts intense, localized energy through cavitation, ^54^ it is unlikely that the SFMIO chains could undergo this dissolution during MPI scans.

This steady-state remanence has implications for the encapsulation of SFMIOs for *in vivo* usage. SFMIOs currently only show superferromagnetism in non-polar solvents, and require packaging to be used in subjects. The long-term remanence of SFMIOs could depend on encapsulation geometry, given its apparent dependence on alignment to the measurement axis and container shape. Optimizing encapsulation chemistry to minimize SFMIO interaction with packaging may simplify remanence behaviour.

To connect remanence and coercivity measurements towards optimal scans, another batch of SFMIOs (characterized in Supplemental Fig. 3: *B*_th_ = 8mT) were scanned in the relaxometer across various frequencies (100Hz - 10kHz) and amplitudes (1 mT - 20 mT). The peak SNR was optimized at high amplitudes, above a frequency threshold (Fig. 5A, at *B*_app_ = 20 mT, *f* ≥ 1 kHz). The sharp SNR drop-off below frequency and amplitude thresholds corroborates slew rate and coercivity thresholds for SFMIOs, as expected from our remanence decay measurements. The SNR optimum at high-amplitude suggests that MPI scan trajectories require a complete rework to best utilize SFMIOs. Given that SFMIO coercivity yields worse scan efficiency at lower amplitudes (Fig 5B), and that MPI safety concerns set frequency limits for a fixed amplitude^55^ (Fig. 5C), it is clear that optimal SPIO scan strategies^35^ (low-amplitude, high-frequency) are unusable for SFMIOs.

**Figure 5:**
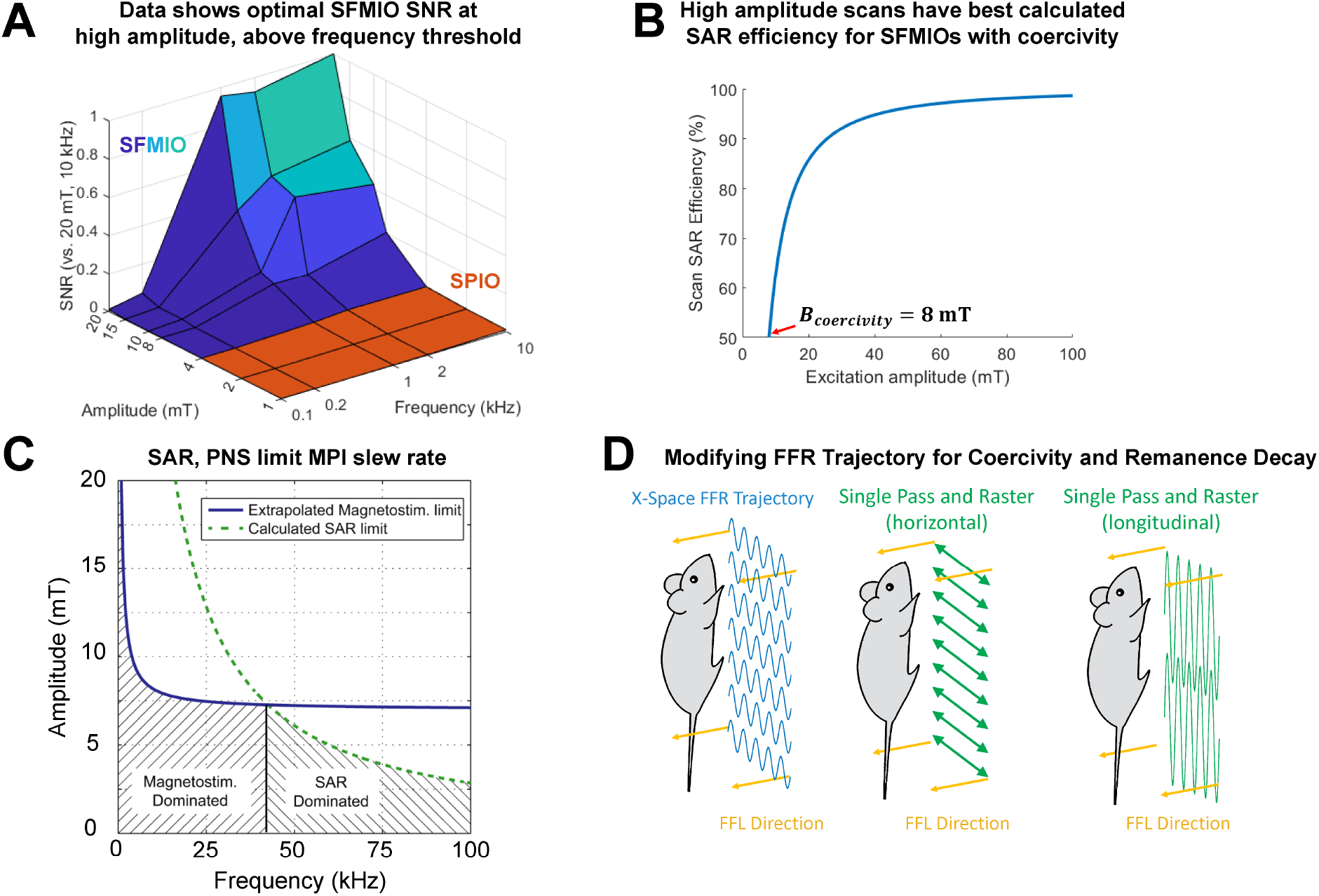
Optimal MPI scanning with remanence decay and coercivity: The coercivity and remanence decay associated with SFMIOs alter what constitutes an optimal MPI scan. (A) A parametric sweep across frequency and amplitude show that optimal SNR efficiency is at high amplitudes (*B*_app_ = 20mT), and above a frequency threshold (*f* ≥ 1 kHz). This sweep reveals the effects of both coercivity and remanence decay, showing extremely low SNR below frequency and amplitude thresholds (B) Coercivity also lowers scan efficiency and unnecessarily heats subjects in lower amplitude scans, pointing towards high amplitude scans. (C) MPI is governed by magnetostimulation and specific absorption ratio (SAR) limits, and requires scan parameters to be under frequency/amplitude thresholds (adapted from Saritas et al. 2013^55^ with permission). (D) Traditional MPI trajectories utilise 20mT, 20 kHz acquisition, running into the aforementioned safety and efficiency concerns, This can be mitigated by increasing scan amplitudes and lowering excitation frequency to the limits specified by remanence decay. Taken to the limit, we propose the Single Pass and Raster (SPaR) sequence, which acquires the entire magnetic field of view in a single half-cycle (*B*_app_ = 99mT for *G* = 6.3Tm^-1^, *FOV* = 30 cm, *B*_coercivity_ = 8mT), and lowers scan frequencies to the limit specified by remanence decay (*f* = 1 kHz). Note that the magnetic field of view may not cover the subject in the axial direction, necessitating mechanical translation.

We propose a new scan that covers the full magnetic field of view per half-cycle: the Single Pass and Raster (SPaR) sequence (Fig. 5D). The frequency would be lowered to accommodate MPI safety limits, down to the limits specified by remanence decay. SPaR could be implemented in the transverse or longitudinal axis, depending on scanner hardware limitations. Future work will focus on SPaR implementation and optimization at higher amplitudes (≈ 99 mT for *FOV* = 30 cm, *G* = 6.3T/m and *B*_coercivity_ = 8 mT).

## Conclusion

This work investigated the effects of SFMIO remanence and coercivity on MPI scanning. We found remanence decay and reformation dependent on applied magnetic fields. Optimal MPI scanning was found to be above minimum excitation frequencies and at high amplitudes, yielding optimal SNR while maintaining their > 10-fold resolution improvement. Remanence decay and coercivity thresholds bound these optimal scan parameters. We propose that further scanner development utilise the Single Pass and Raster sequence, using high-amplitude, low-frequency excitation to safely and efficiently scan SFMIOs. The resolution improvements from SFMIOs could lead to 100-fold cheaper scanners, and are crucial for the clinical translation of MPI.

## Acknowledgement

The authors thank: the Salahuddin Lab (especially JCH Hsu) at UC Berkeley for usage of their VSM; the Electron Microscopy Lab at UC Berkeley for usage of their equipment; ZW Tay, PhD, XY Zhou, PhD, and QL Huynh for discussions.

## Author Contributions

KLBF and SMC conceptualized remanence models and initial remanence experiments. KLBF developed, executed, and analyzed the followup experiments. JB carried out sonication experiments. KLBF and CC acquired the SFMIO images. RK performed iron quantification. KLBF, CS, and YL assessed remanence models. CC, CS, YL, BDF, PC, and SMC refined methodology. JB, JML, AH, and KY synthesized SFMIOs. KLBF prepared the draft and figures. All authors reviewed and approved the final draft.

## Funding

The authors acknowledge support from: NIH grants R01s EB024578, EB029822, T32 GM 098218 and R44 EB029877, UC TRDRP grant 26IP-0049, UC Discovery Award, M. Cook Chair, Bakar Fellowship, NSERC PGSD3-532656-2019 Fellowship, UCB Bioengineering Craven Fellowship, UC CRCC Doctoral Fellowship, Siebel Scholars program, NSF GRFP.

## Conflict of interest

SMC is a co-founder of an MPI company, Magnetic Insight, and holds stock in it. The authors declare no other conflicts.

## Supporting Information Available

**Supplemental Figure 1**: MPI images of the phantoms in Fig 1A-C, normalized to peak intraimage signal.

**Supplemental Figure 2**: Remanence decay vs. *B_SS_* for *M_SFMIO_* > 0 (complementary to Fig. 3C).

**Supplemental Figure 3**: Slew-normalized PSFs for Fig. 5A’s SFMIOs and theoretical coercivity losses in MPI scans.

## TOC Graphic

Superferromagnetic iron oxide nanoparticles (SFMIOs) have shown 10x resolution boosts in magnetic particle imaging due to interacting nanosparticles. We show SFMIO remanence decay, reformation, and frequency thresholds in MPI scans, suggesting high amplitude, low frequency scans for optimal *in vivo* SFMIO usage. SFMIOs will provide a 10× scanner field strength reduction and thus up to 100× cost reduction.

## Notes

### Competing Interest Statement

SMC is the co-founder of an MPI company and hold stock in that company. The authors declare no other interests.

