## Supplemental Figures for "First superferromagnetic remanence characterization and scan optimization for super-resolution Magnetic Particle Imaging"

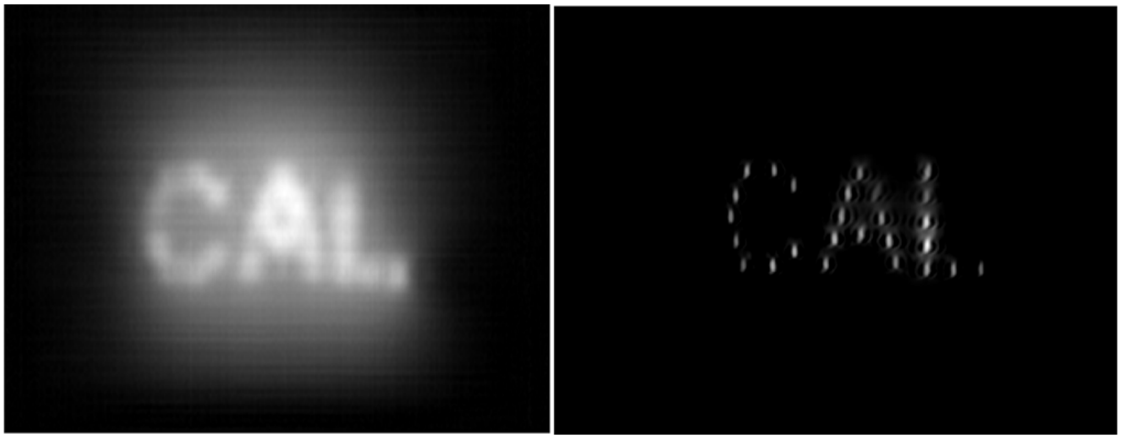

Figure 1: The MPI images of ferucarbotran (VivoTrax$^{TM}$, left) and SFMIO (right) phantoms from Fig 1A-C, normalized to peak intraimage signal (i.e. $\max(signal(x, y)) = 1$ for both images) to clearly show resolution. Note the significantly worse resolution in the image on the left.

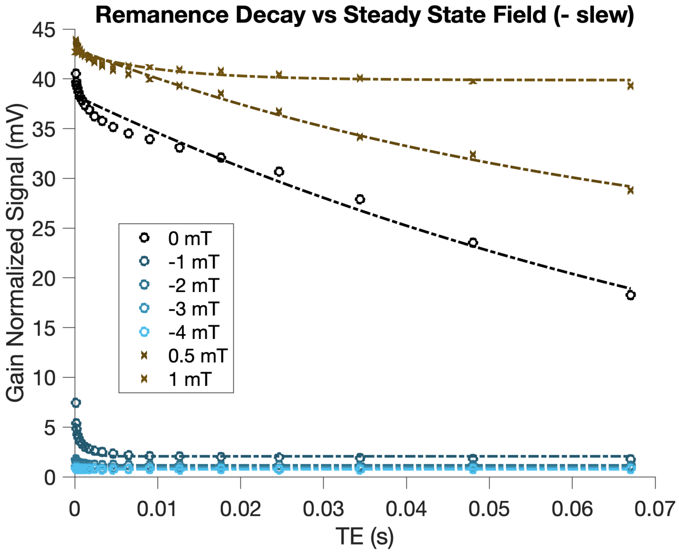


Figure 2: The remanence decay in the opposite slew direction versus Fig. 3C. Note the rapidly increasing decay, where measurements for bias fields stronger than -2 mT show remanence decay faster than can be measured (i.e. $\tau_{decay,|B|>2\,\mathrm{mT}} < 100\,\mu\mathrm{s}$

.

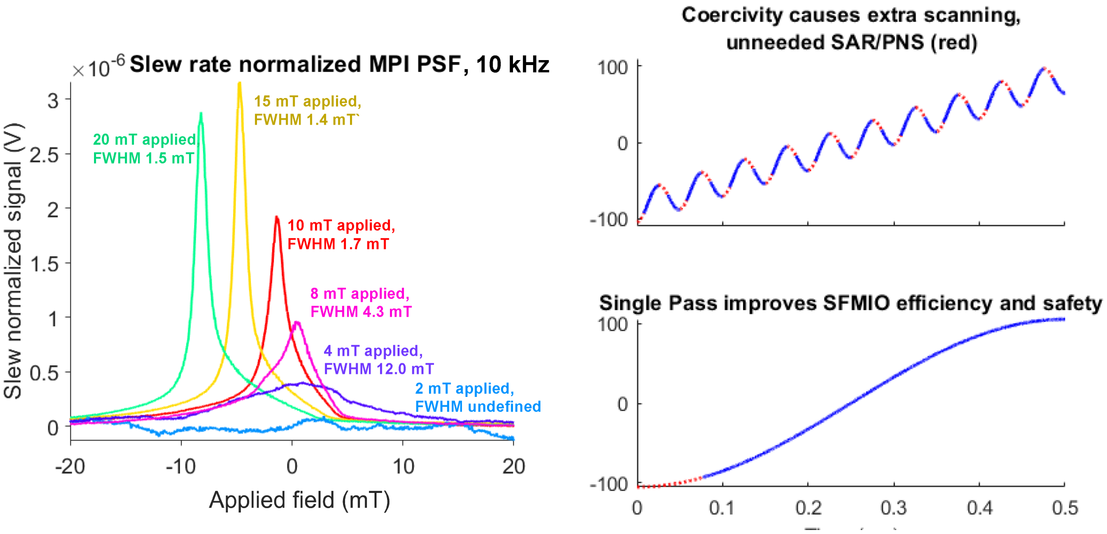


Figure 3: **Left**: the slew-normalized characteristic PSFs for Fig. 5A's SFMIOs, showing the SFMIO amplitude threshold of 8 mT. Note how despite slew normalization, the signal increases with amplitude, which does *not* happen in SPIOs. **Right:** a brief diagram showing wasted percentages of scans due to coercivity (8 mT in this diagram). This results in the wasted SAR as shown in Fig. 5B, and shows why high amplitude, low frequency scans are preferable for SFMIOs
